# Unifying intra- and inter-specific variation in tropical tree mortality

**DOI:** 10.1101/228361

**Authors:** James S Camac, Richard Condit, Richard G FitzJohn, Lachlan McCalman, Daniel Steinberg, Mark Westoby, S Joseph Wright, Daniel S Falster

## Abstract

Tree death is a fundamental process driving population dynamics, nutrient cycling, and evolution within plant communities. While past research has identified factors influencing tree mortality across a variety of scales, these distinct drivers are yet to be integrated within a unified predictive framework. In this study, we use a cross-validated Bayesian framework coupled with classic survival analysis techniques to derive instantaneous mortality functions for 203 tropical rainforest tree species at Barro Colorado Island (BCI) Panama. Specifically, we develop mortality functions that not only integrate individual, species, and temporal effects, but also partition the contributions of growth-dependent and growth-independent effects on the overall instantaneous mortality rate. We show that functions that separate mortality rates into growth-dependent and growth-independent hazards, use stem diameter growth rather than basal-area growth, and attribute the effect of wood density to growth-independent mortality outperform alternative formulations. Moreover, we show that the effect of wood density – a prominent trait known to influence tree mortality – explains only 22% of the total variability observed among species. Lastly, our analysis show that growth-dependent processes are the predominant contributor to rates of tree mortality at BCI. Combined, this study provides a framework for predicting individual-level mortality in highly diverse tropical forests. It also highlights how little we know about the causes of species-level and temporal plot-scale effects needed to effectively predict tree mortality.

---

Rates of plant mortality are known to vary widely among individuals within species, among coexisting species, between forests, and from year-to-year [1–3]. This variation has considerable consequences for forest structure and dynamics. For example, death of a single large tree can transfer up to 20000 kg of carbon from living to decaying carbon pools [4]. Furthermore it creates a gap in the canopy that can restart a successional race, during which 100’s of plants may die while competing for a spot in the sun. In models of forest dynamics, variation in mortality rates has been shown to have a larger impact on forest structure than variation in absolute growth rates [5]. Improving our understanding of the mortality process is therefore a priority for making accurate predictions about population, carbon and nutrient dynamics of forests; especially in an era of rapid environmental change.

Two difficulties arise when studying tree mortality in tropical rainforests. The first is that large population sizes and long periods of observation are required to make inferences into how various factors affect mortality rates [3]. This requirement arises in part because the observable outcome – alive vs. dead – is one step removed from the variable we ideally want to measure: λ*_i_*(*t*), the instantaneous rate of mortality for individual *i* at time *t*, also called the “hazard function”. In classic survival analysis [6], the probability an individual dies between times *t*_2_ and *t*_3_ is a function of λ*_i_*(*t*):

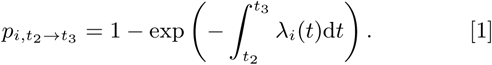

Here 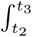 λ*_i_*(*t*)dt is the “cumulative hazard” between *t*_2_ and *t*_3_ [6]. The observed survival outcome *S*_*i*,*t*_2_→*t*_3__ (0 = alive, 1 = died) is then a realisation of this probability

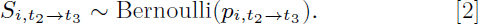

As trees are long-lived and we are trying to estimate the shape of a continuous hazard function (λ*_i_*(*t*)) from binary data, large sample sizes are required. Detailed studies of tree mortality have thus only recently become possible, with the accumulation of growth and survival data from repeat surveys spanning several decades in plots containing thousands of individuals [7].

Another difficulty when studying mortality – similar to that faced in other fields like medicine and engineering – is determining the shape of hazard functions that skilfully predict patterns in mortality. Past research has identified a range of factors with significant effects on tree survival, including an individual’s growth rate [1, 2], traits [3, 8–10], and size [11]. However, these influences have not yet been integrated into a common hazard function [12–14].

### Significance Statement

Tree mortality is a fundamental demographic process affecting forest dynamics and carbon cycling. Here, for the first time, we use over 400,000 observed survival records collected over a 15 year period from more than 180,000 individuals, to simultaneously estimate growth-dependent and growth-independent mortality across 203 tropical forest tree species. We found that growth-dependent mortality was the predominate factor influencing tree mortality rates at Barro Colorado Island. Furthermore, we found that while wood density influenced mortality rates by decreasing growth-independent mortality, wood density only accounted for a small fraction of the overall species variability in mortality rates, suggesting that there must be other species traits that strongly affect mortality.

## Towards a unified model of tree mortality

A specific challenge in developing hazard functions for plants is to estimate the relative contribution of growth-dependent and growth-independent hazards on an individual’s overall hazard. While plants die via many causes, these broadly fall into two categories. The first are growth-dependent hazards, where plants die because of insufficient carbon assimilation for growth and repair. The second are growth-independent hazards, where plants die because of stochastic events, irrespective of their growth rate, such as windfall or fire. An individual’s total hazard is the sum of growth-independent and growth-dependent components.

A further challenge in developing hazard functions for plants is to integrate the conflicting relationships observed within and across species between mortality and growth. Within species, mortality rates are lower for fast-growing individuals, presumably because those individuals have superior carbon budgets and are thus able to tolerate or repair diverse stresses [1, 12, 15]. Empirical studies broadly support this theory, with many indicating exponential declines in mortality with increased growth rate, *X_i_*(*t*) [1, 11, 16]. By contrast, across species, there is a strong trade-off between growth and survival, with individuals from faster-growing species exhibiting higher mortality rates than individuals from slower-growing species [3, 9]. This trade-off may arise via traits with antagonistic effects, such as wood density. Denser wood – which is expensive to build and thus slows growth – reduces stem breakage [17, 18], embolism [19, 20], pathogen attack [21], and thereby mortality [3, 8, 9, 18].

Here, we attempt to reconcile intra- and inter-specific factors as well as partition instantaneous mortality rates into growth-dependent and growth-independent rates. We achieve this by evaluating the following, unified hazard function, incorporating individual-, species-, and census-level effects:

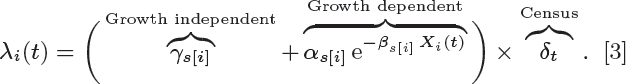

Eq. 3 allows for additive growth-independent and growth-dependent effects, includes a negative exponential effect of growth rate, and allows for mortality to vary among censuses, via the random effect *δ_t_*. Further, the parameters *α_s_*, *β_s_* and *γ_s_* vary by species, *s*. Here we include an effect of a species trait (wood density, *ρ*), as well as a species random effect that captures any remaining species-level differences not accounted for by wood density (for details see Methods).

To validate this model, we fit a series of models with increasing complexity (eq. 3 being the most complex) and compare their skill in predicting patterns of tropical tree mortality for 180,509 individuals from 203 tree species at Barro Colorado Island (BCI), Panama (Fig. 1). The data are repeat censuses of stem diameter and tree status (alive vs. dead) taken over a 15 yr period (Fig. 1A). In total 427,468 observations were used to fit these models. We compare the skill of different hazard functions in predicting outcomes in novel data (i.e. not used in model fitting) via 10-fold cross-validation (Fig. 1D) [22, 23]. Evaluating models in this way is computationally expensive, making it impossible to run all possible model formulations. We therefore fit models across five iterative stages of model development. At each stage, the best model from the previous stage was taken as input for the next stage.

**Fig. 1.**
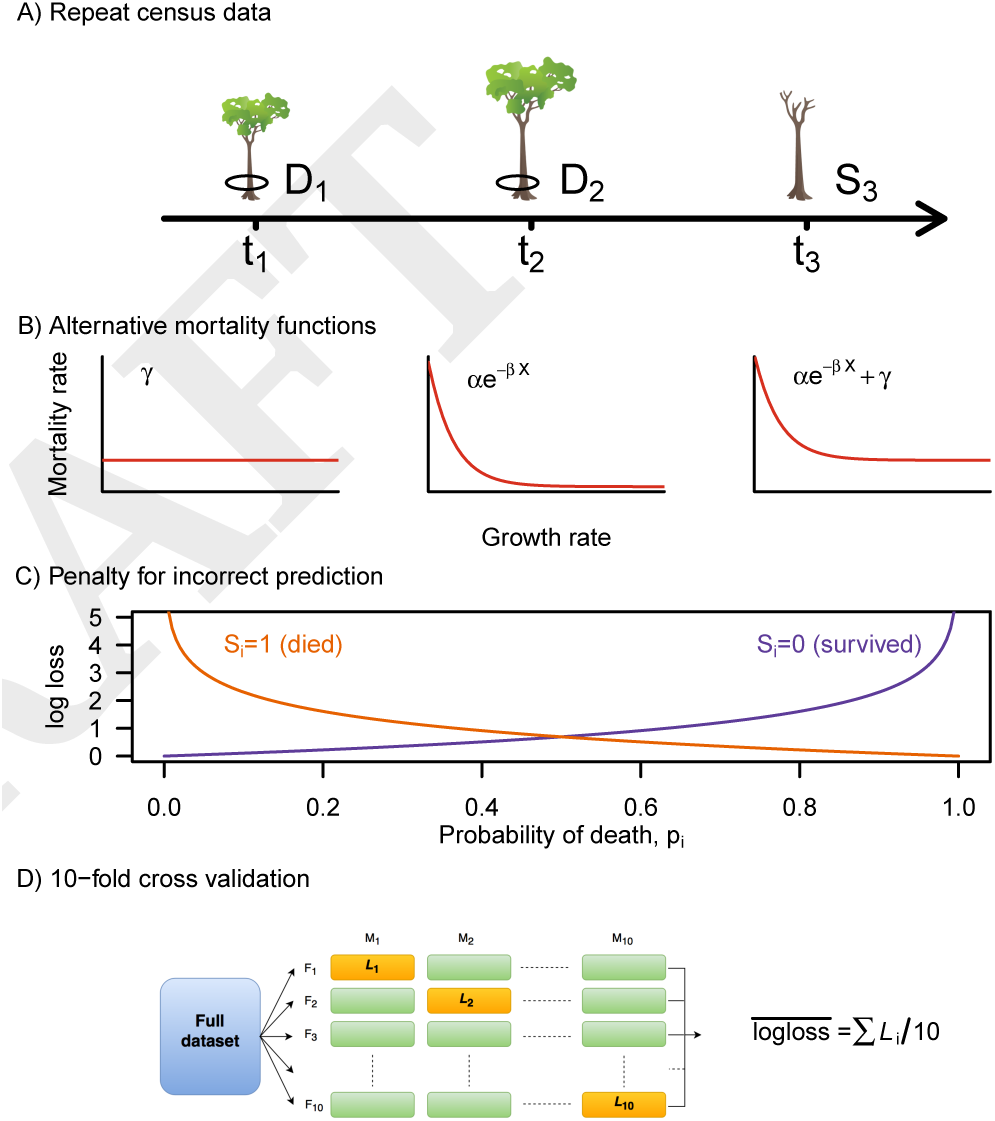
Outline of methodology. A) Our data consist of repeat measures of stem diameter (*D*) and status (*S*, 0 = alive or 1 = dead) for individual trees at specific census dates (*t*_1_, *t*_2_, *t*_3_). B) We consider three alternative hazard functions: 1) a baseline hazard, 2) a growth-dependent hazard; and 3) a function that combines both baseline and growth-dependent hazards. The parameters of the models are biologically interpretable: *α* defines the instantaneous mortality rate at low growth rate; *β* reflects the sensitivity of mortality rate to changes in growth rate; and *γ* is the asymptote, or baseline hazard. Combined *α* and *β* capture growth-dependent mortality, while *γ* captures growth-independent mortality (e.g. windfall, fire) that kill a plant, irrespective of its growth rate. For each model form, we consider two alternative predictors of growth, *X* (basal area and stem diameter growth), as well as allowing for species-level effects on the parameters *α*, *β* and *γ*. C) Each model’s skill in predicting observed outcomes (*S*) is quantified via the log-loss function (eq. 5). D) The predictive skill of alternative models was evaluated via 10-fold cross validation. The entire dataset is split into 10 folds (*F*_1_, …, *F*_10_). Alternative models were fit 10 times (*M*_1_, …, *M*_10_), using different combinations of testing (1 fold; orange) and training (9 folds; green) data. Predictive capacity was assessed by averaging the log loss’s obtained from the 10 test data predictions.

In stage 1, we ask whether mortality rates vary substantially between censuses, to establish whether a census effect is required. The null model is a constant, invariant with respect to species, growth rate, or year. In stage 2, we assess whether a hazard function including both growth-dependent and growth-independent terms outperform a function including only one of these. Species effects were excluded, so we are simply asking which of three functional forms for λ*_i_*(*t*) (Fig. 1B) best predicts the data. The simplest form (Fig. 1B, left) assumes a constant growth-independent hazard rate. The second form (Fig. 1B, middle) assumes the risk of dying declines towards an asymptote of zero as growth increases. The third form (eq. Fig. 1B, right) is the summation of thetwo previous models; towards an asymptote of zero as growth increases. The third form (eq. Fig. 1B, right) is the summation of the two previous models; this allows λ*_i_*(*t*) to decrease exponentially with increasing growth rate, but, unlike a standard negative exponential, asymptoting at some baseline hazard > 0. Combined, these three models capture a variety of functional responses previously proposed, including effects represented in current vegetation models [14, 24], which have not previously been systematically compared. We also investigate which growth measure (stem diameter, stem area increment) more skilfully predicts growth-dependent mortality. In stage 3, we examine whether including wood density (a species-level trait) improved model skill, and if so, whether the effect of wood density was on growth-dependent, growth-independent, or both hazards. Finally, we fit a model that allowed parameters to vary by species-level variation that was otherwise not captured by wood density. This allowed us to ask what proportion of interspecific hazard variability is explained by wood density. Using the final “best” model we also conducted post-hoc tests, to determine how species-level parameters were associated with their maximum size and light requirement.

## Results

### Mortality over time

Comparing the three 5-year intervals between censuses from 1995 to 2010, we found average mortality rates progressively decreasing over time. The highest proportion of trees death occurred between 1995–2000 (16%) followed by 2000–2005 (13%) and 2005–2010 (12%). Consequently, when we allowed hazard rates to be scaled by census (i.e. adding term *δ*) we observed a small, but significant, increase in predictive skill (Fig. 2A). Individual census effects can be found in Table S1.

### Hazard functions

Comparing the three hazard functions in Fig. 1B, we found that the third function — with both growth-dependent and growth-independent terms, i.e. (*αe*^−*βX_i_*^ + *γ*)*δ_t_* — significantly outperformed both the growth-independent only (i.e. null) or growth-dependent only functions (Fig. 2A). Moreover, we found that predictive skill was higher when using stem diameter growth over stem-area growth (Fig. 2A). A summary of hyper parameter estimates can be found in Table S1.

### Wood density and other species effects

Including wood density as an effect on either growth-independent or growth-dependent parameters significantly improved model skill relative to a model without such effects (Fig. 2 A,B). The most parsimonious model, with highest predictive skill, was that attributing the wood density effect to the growth-independent hazard term (*γ*; Fig. 2 B). Specifically, we found that wood density was negatively correlated with a species’ baseline mortality rate, *γ* (Fig. 3). This meant that fast growing individuals from a low wood density species had, on average, higher mortality rates (Fig. 4 A,B), and thus higher probability of death across a 1-yr period (Fig. 4 C,D), relative to fast growing individuals of high wood density species. For example, a species with a wood density of 0.3 g cm^−3^ had an estimated mean probability of dying of 0.05 yr^−1^ compared to 0.01 yr^−1^ for a species with a wood density of 0.8 g cm^−3^. Incorporating an additional species-level random effect to capture any additional inter specific differences substantially improved model skill relative to a model with only wood density (Fig. 2A). See Figs. S3-S5 for species-level parameter estimates.

**Fig. 2.**
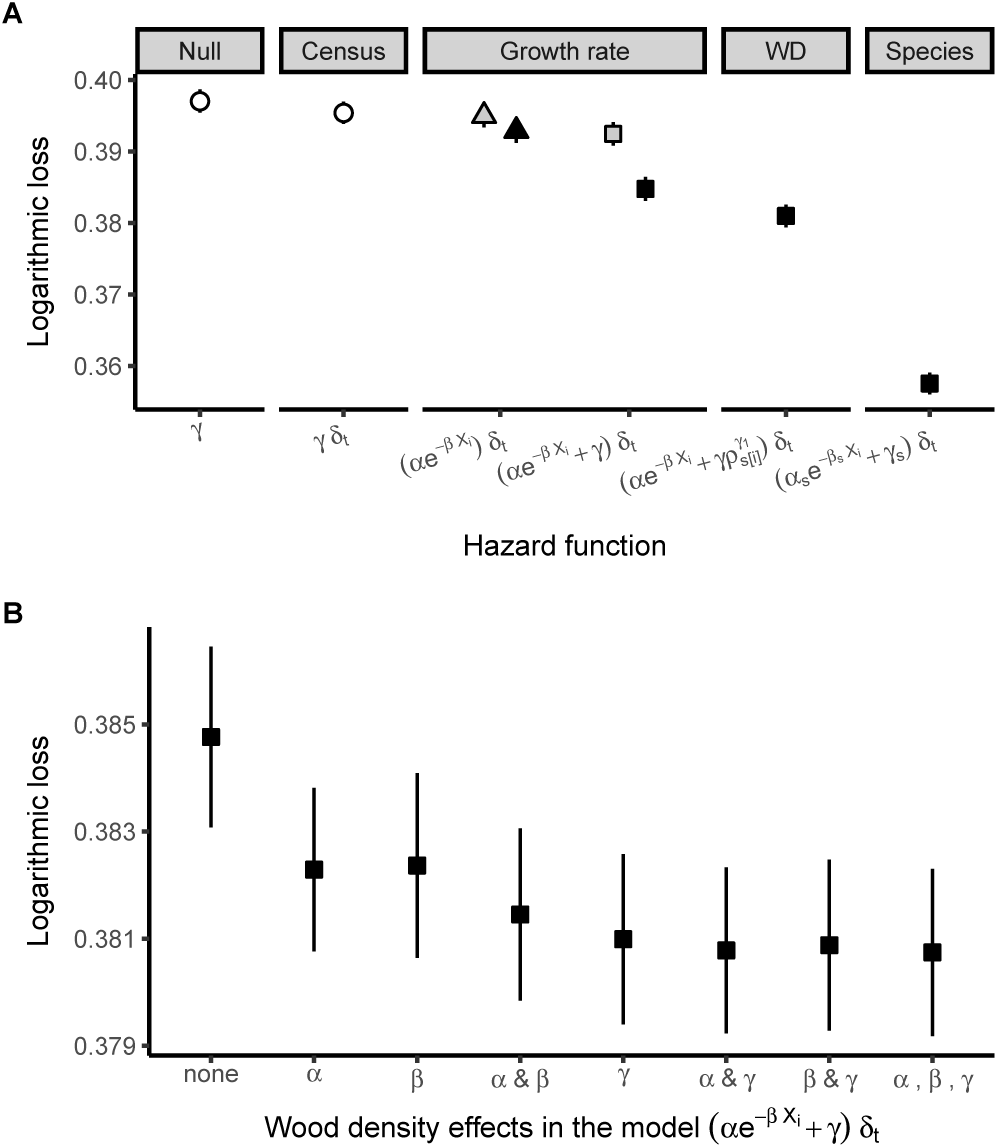
Predictive skill of alternative hazard functions. Values are mean (± 95% credible intervals) logarithmic loss, with lower values implying greater predictive skill. A) Five sequential stages of model selection with each stage increasing in complexity. Null: constant (*γ*) Census: inclusion of census effects (*δ_t_*); Growth rate: the inclusion of growth rate. This includes two possible hazard functions: growth-dependent only hazard ((*αe*^−*βX_i_*^)*δ_t_*) and baseline + growth-dependent hazard with census effects ((*αe*^−*βX_i_*^ + *γ*)*δ_t_*); WD: inclusion of wood density effect (*ρ*) on *γ*; Species: the inclusion of species random effects *s* on all parameters. B) Predictive skill for wood density parameter combinations. Shading represent growth measure used: white = no growth measure, grey = basal area growth, black = dbh growth. For both panels symbols represent functional form: null (circle), growth-dependent hazard (triangle), both growth-independent and growth-dependent hazards (square).

**Fig. 3.**
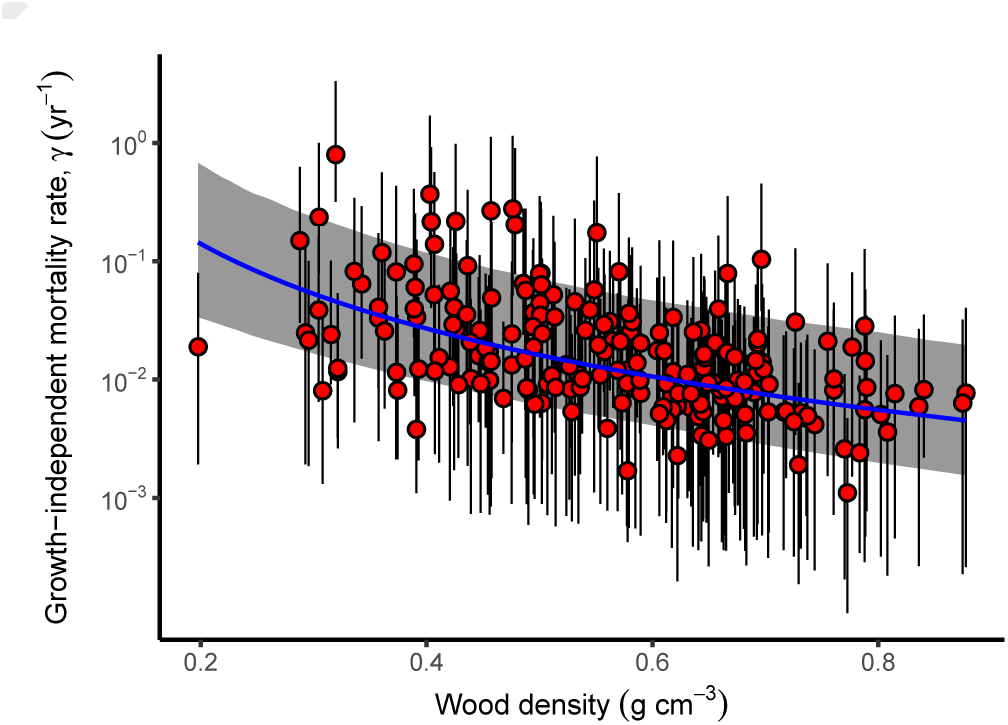
Relationship between species growth-independent mortality rate and wood density. Points are estimated mean baseline mortality rates ± 95% credible intervals for each of the 203 BCI species used in this study. Blue trendline with grey shading shows average (± 95% credible intervals) expected relationship.

**Fig. 4.**
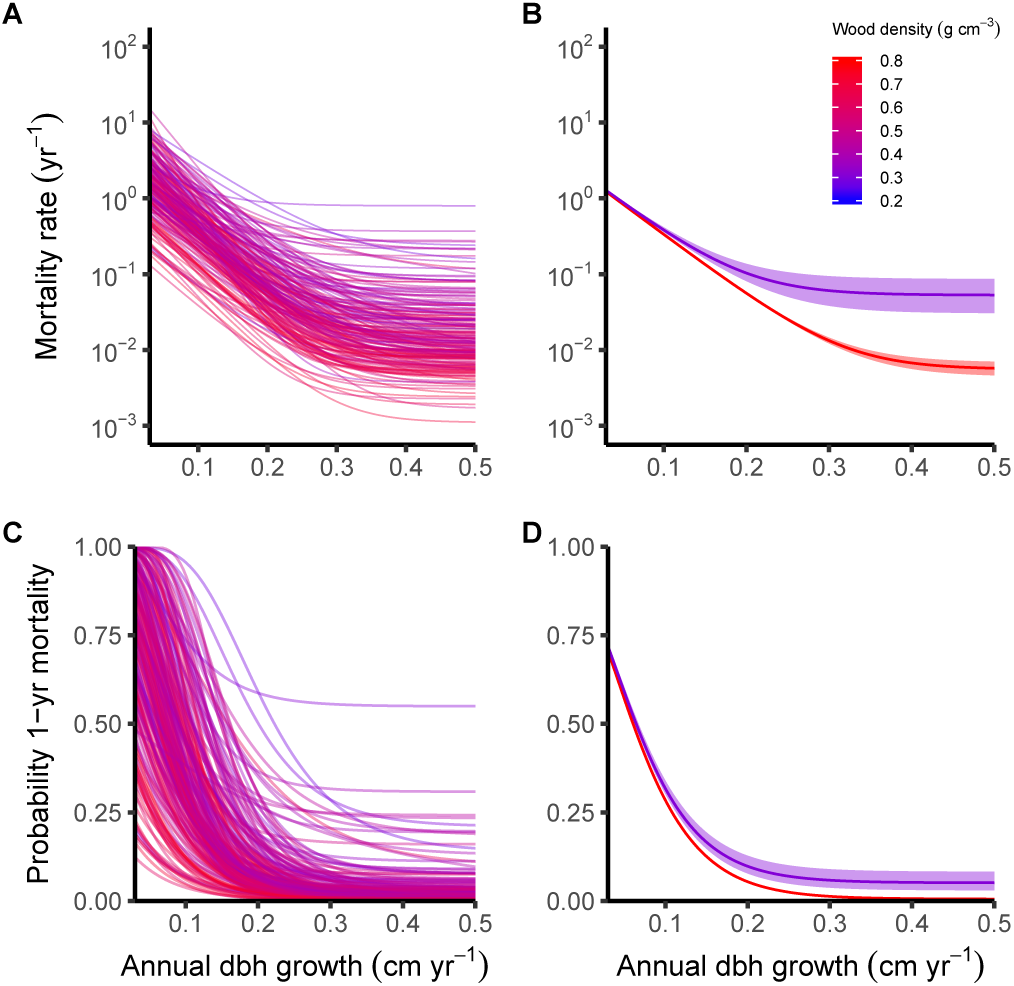
Predicted changes in instantaneous mortality rate (A,B), and 1-yr mortality probability (C,D), with increasing growth rate. A,C) Estimated curves for all 203 BCI species used in this study. B,D) The average (± 95% credible intervals) curve for a high (0.8) and low (0.3) wood density species.

### Parameter contribution to predicted variation

To compliment the analysis, we used the final fitted model to estimate the amount of variation captured by different factors in the predicted probability of dying across 1 year for all individuals (Fig. 5). We achieved this by removing each parameter and calculating the reduction in the sum of squares for this model relative to the full model. Species effects contributed 80% of predicted variation in 1-yr mortality. Wood density explained 22% of this species variation. Census accounted for 6.1% of the total predicted variation. The growth-dependent hazard accounted for 68% of the total variation, while growth-independent hazards accounted for the remainder (Fig. 5).

### Mortality rates, maximum species size and species light requirements

To determine whether species-level parameters were associated with their maximum size or light requirement, we conducted a post-hoc analysis comparing the fitted parameters to these traits. We found that both *α*, the effect of low growth rate, and *γ*, the growth-independent hazard term, were weakly and positively correlated with gap index (a measure of species light requirement; Fig. 6 A,C,E). By contrast, *β* (the parameter that defines the exponential decay with increasing growth rate) was negatively correlated gap index. These correlations suggest that species which predominately recruit in gaps are more prone to death, due to both low growth and stochastic chance. They also required faster growth rates to achieve a given mortality rate relative to a species that recruit in shade. Correlations between estimated species parameters and their associated maximum DBH were weak (Fig. 6).

**Fig. 5.**
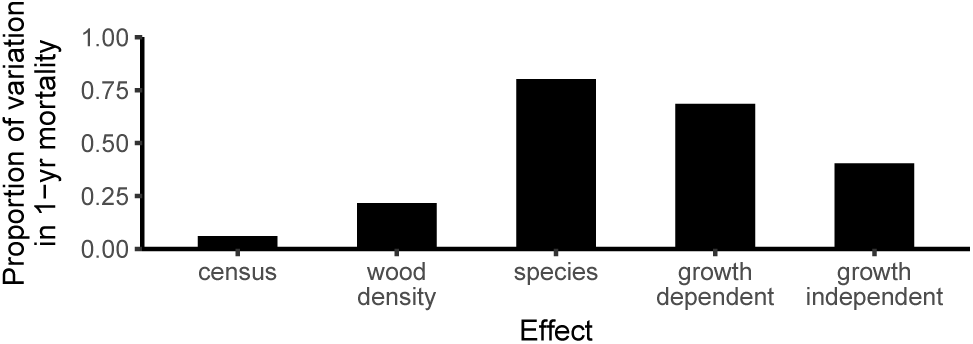
Proportion of variation in predicted 1-year mortality for all individuals captured by different effects. Effects are not mutually exclusive thus sum to more than 1.0. Note “wood density” is a subset of the overall “species” effect.

**Fig. 6.**
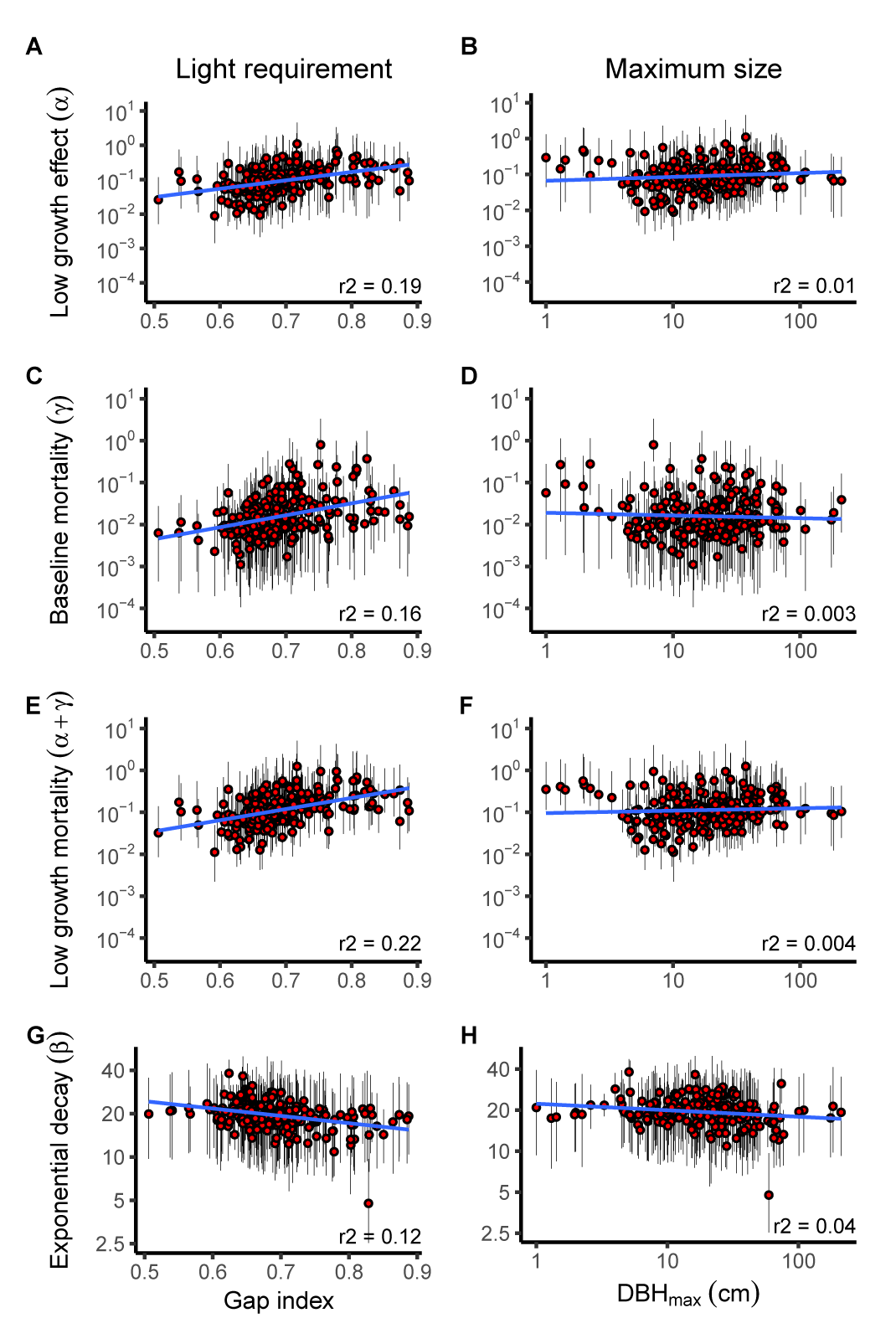
Posthoc correlations between estimated species parameters and measures of species light requirement (left) and species maximum DBH (right). Points are means (± 95% credible intervals) for each of the 203 species used in this study. Blue line shows the average trendline as determined from standard linear regression. Gap index is a species level index of the light required for recruitment. Low gap index indicates the species readily recruits under dense canopy and 1 indicates that a species only ever recruits in gaps.

## Discussion

Our Bayesian framework coupled with cross validation revealed that the most explanatory and parsimonious model of tropical tree death was that which: 1) partitioned mortality into growth-dependent and growth-independent hazards; 2) used stem diameter growth rather than basal-area growth; 3) attributed the effect of wood density to growth-independent mortality; and 4) incorporated temporal variability. Moreover, we found that rates of tropical tree mortality varied substantially between species and that wood density, a species level functional trait, explained only a limited proportion of the overall inter-specific variation.

The findings of this study provide empirical support for dynamic vegetation models that estimate mortality as the sum of growth-dependent and growth-independent hazards [14, 24, 25]. We show that regardless of growth measure, incorporating both hazards significantly improves model predictive skill. This is because the growth-dependent hazard allows for deaths associated with low carbon budgets, and as a consequence, incorporates intra-specific variability attributed to carbon related stresses (e.g. competition, parasites, herbivory). By contrast, the growth-independent hazard accounts for deaths caused by events that arise irrespective of an individual’s growth rate (e.g. windthrow, lightning strike).

Additionally, the partitioning of mortality into growth-dependent and growth-independent effects allowed us to estimate the proportion of variation attributed to each. Like many other studies [26–28], our analyses highlight the importance of light competition in influencing tropical tree demographic rates. Specifically, we found that the growth-dependent hazard accounted for 68% of the total predicted variability in mortality rates (Fig. 5). This suggests deficiencies in carbon budget are a major contributor to tree death on BCI.

Incorporating the effect of wood density on mortality rates also improved predictive performance. Our analyses revealed that the most parsimonious combination of wood density effects was when it was attributed to only the growth-independent hazard term. Specifically, high wood density species had lower baseline rates relative to low wood density counterparts. This finding corroborates the observed negative correlation observed between mortality and wood density reported elsewhere [8, 18]. More importantly, our analyses support the theory that wood density reduces mortality rates by decreasing a species’ vulnerability to growth-independent threats, such as windthrow, trampling and treefall [17, 18].

While wood density effects are now being incorporated in mortality algorithms of many vegetation models [24, 29], our analysis indicate that such effects are likely to only capture a small proportion of the overall inter-specific variation (Fig. 4–5). This means that such models are likely to severely under-estimate the true variability in mortality rates. Consequently, this underestimation is likely to manifest in biased estimates of carbon, water and nutrient dynamics of ecosystems [5].

We found that wood density only explained 22% of the total 80% variation explained by species effects, suggesting that other traits are also affecting tree mortality rates. Post-hoc analyses suggest this unexplained variation was in part related to a species’ light demand, but not its maximum height. Both traits have been proposed major axes of inter-specific variation in tropical rainforests [30] (Fig. 6). Specifically, light demanding species (i.e. high gap index) had higher growth-independent hazard rates and were more susceptible to dying as a result of low growth, relative to those that readily recruit in shade (i.e. low gap index), supporting past findings [27]. By contrast, we detected no correlations between a species maximum stem diameter and mortality rates, both contradicting [30] and supporting [9] previous results. Future research should therefore resolve how other traits influence growth-dependent and growth-independent hazards, and identify the combination of traits needed to improve model predictive skill.

Future research should also resolve how climate influences both growth-dependent and growth-independent hazards, particularly as climate-driven mortality is increasing [31, 32]. By contrast, our analyses revealed a marginal decline in mortality at BCI from 1995 to 2010. Attributing temporal changes to particular climatic variables is challenging, however, due to low number temporal replication (n=3).

We should also consider how the interval between censuses affects the estimation of growth-dependent and growth-independent hazards. Large census intervals may underestimate growth-dependent mortality, as wide census intervals will
not capture deaths due to rapid declines in growth, or events such as drought (although drought might also increase tree growth [33]). Consequently, we may overestimate the relative contribution of growth-independent hazards.

Here we showcase a new framework for modelling tropical tree mortality that unifies empirical evidence from within and between species studies. This framework also provides an approach for partitioning mortality rates into growth-dependent and growth-independent hazards. Our findings reveal that while wood density is an important trait affecting mortality rates, we are still only capturing a fraction of the overall species variability in mortality rates.

## Materials and Methods

### Data

We derived plant mortality models using individual growth and survival data collected from a relatively undisturbed 50-ha tropical rainforest plot on BCI, Panama (9.15°;N, 79.85°;W). The climate on the island is warm and rainfall is seasonal with most falling between April and November [34].

Within the 50-ha plot the diameter at breast height and survival status of all free-standing woody plants that were at least 1.3 m tall and had diameter ≥ 1 cm were recorded in 1981—1983, 1985, and every 5 years thereafter [34]. For the purpose of modelling mortality as a function of past growth, we discarded data collected prior to 1990. This was because diameter measurements were rounded to the nearest 5 mm for individuals with dbh < 55 mm, whereas in later censuses all individuals were measured to the nearest millimetre [33]. Consequently, we modelled tree mortality as a function of past growth for censuses 1995–2000, 2000–2005 and 2005–2010. We discarded species that do not exhibit secondary growth (e.g. palms and ferns), contained fewer than 10 individuals or did not contain an estimate of wood density. We also excluded individuals that: 1) did not survive at least two censuses (two being required to estimate growth rate); 2) were not consistently measured at 1.3 m above ground; 3) were multi-stemmed; 4) resprouted or seemingly “returned from the dead”; or 5) were extreme outliers – stems which grew more than 5 cm yr^−1^ or shrunk more than 25% of their initial diameter. In total 427,468 observations were used in this study comprising 180,509 individual trees and 203 species. Because of computational costs, the models fit in this study do not include individual random effects, as this would require estimation of an additional 180,509 parameters. Instead, our models assume that repeat measurements of an individual are independent of one another. We believe this is a reasonable assumption given that there is approximately 5-years between censuses.

Wood density for each species was estimated by coring trees located within 15 km of the BCI plot [9]. Cores were broken into pieces, each 5 cm long and specific gravity of each piece was determined by oven drying (100°C) and dividing by the fresh volume (as measured by water displacement).

### Model fitting

Eqs. 1-3 were fit to the data using Bayesian inference and with covariates for growth rate in previous census and wood density, as well as random effects. Growth rates were estimated from field measurements of diameter, which inevitably include observation error. In our dataset, 8% of estimated growth rates were negative. To ensure our mortality model was not biased by these unlikely values we first applied a probabilistic model to estimate *“true growth”,* taking into account measurement error and the distribution of growth rate across the community (see Supplementary Material S1 for details; Fig. S1). The parameters as, *α_s_*, *β_s_* and *γ_s_* were modelled as a function of both wood density (measured at species level) and a species-level random effect:

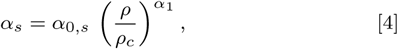

with similar formulations for *β_s_* and *γ_s_*. Here *α*_1_ captures the effect of wood density *ρ* on *α_s_*, while *α*_0,*s*_ captures any other species-level residual error not explained by wood density for species s. These random effects were modelled as random realisations from log-normal distributions. The form of eq. 4 ensures that parameters remain positive; and on a log scale this equates to an additive linear model centered around *ρ_c_*. We also centered growth rate *X_i_* at the lower 5% quantile for both diameter increment and area growth (0.172 and 0.338, respectively), meaning *α_s_* should be interpreted as the hazard rate when growth rate was very low. Weak priors on all hyper-parameters were set (see Supplementary Material S2 for details). Models were fit in R 3.4.1 using the package `rstan` 2.16.2 [35] and employing some numerical optimisations (see Supplementary Material S3-S4 for details). We executed three independent chains and in all cases modelled parameters converged within 2000 iterations. Convergence was assessed through both visual inspection of chains and reference to the Brooks-Gelman-Rubin convergence diagnostic [36]. After discarding the first 2000 iterations as ‘burn in’, a further 2000 iterations were taken from the joint posterior. Species parameter estimates from the final model are shown in Figs. S3–S5.

### Evaluating model skill

Predictive skill was quantified by estimating the average log loss across 10-folds for held-out data, 𝓛̅ (Fig. 1). Logarithmic loss – commonly known as log loss, 𝓛, measures the skill of a model by penalizing incorrect predictions, based on how wrong the predicted probability is from the observed outcome, *S_i_* (Fig. 1C). Lower 𝓛 implies greater skill. The average log loss across all individuals for the fcth fold of held-out data, 𝓛*_k_*, is then

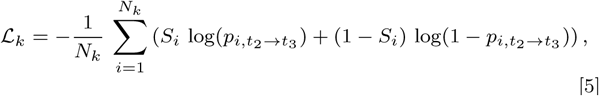

where *Nk* refers to the number of observations in fold.

### Posthoc correlations

We calculated a gap index as a measure of a species’ light dependence using annual canopy census data collected during 1985–1990 and 1990–1995. The canopy census recorded, in all 5 by 5 m subplots across the 50 ha plot, the presence of leaf in six height intervals (0-2,2-5,5-10,10-20,20-30, >30 m). For each subplot, we calculated the number of strata > 2 m containing vegetation; and then transformed this to a light index ranging from 0 (dense shade) to 1 (gap). As light may penetrate into a subplot from the edge of a subplot, we rescaled this index to account for values in the eight immediate neighbouring subplots. Specifically, we used a weighted sum approach whereby the central subplot is assigned a weight of 8 and the eight neighbouring subplots are assigned a weight of 1. This meant that the contribution of the central plot was equivalent to the combined effect of all eight neighbouring plots. These weighted values were then summed and rescaled between 0 and 1 by dividing by the maximum value estimated across all subplots. The gap index for each species was estimated as the mean light index encountered by new saplings appearing in the census (Fig. S2).

## ACKNOWLEDGMENTS

We thank H. Muller-Landau and anonymous reviewers for feedback and B. Carpenter for technical advice. We acknowledge S. Hubbell, R. Foster, R. Pérez, S. Aguilar, S. Lao, S. Dolins, & hundreds of field workers for their contribution; and the National Science Foundation, Smithsonian Tropical Research Institute, & MacArthur Foundation for funding the design, collection, quality control and management of long-term growth data at BCI. Fig. 1a uses images by Tracey Saxby, IAN Image Library. J.C., R.F., L.M. and D.S. were supported by the Science and Industry Endowment Fund (SIEF; RP04-174). D.F. and M.W. were supported by fellowships from the Australian Research Council.

## Notes

The authors declare that they have no competing financial interests.

